# HIV-1 virions selectively package circESYT2 to sculpt an actin scaffold that constraints egress

**DOI:** 10.64898/2026.01.21.700838

**Authors:** Pratibha Madbhagat, Aniruddha Panja, Ajit Chande

## Abstract

Retroviruses such as HIV-1 package both viral and host RNAs, yet whether circular RNAs (circRNAs) enter virions has remained unknown. Here, we capture the HIV-1 RNA packageome and show the evidence that circRNAs represent a previously unrecognized class of selectively encapsidated host RNAs. Using nanopore sequencing of purified virions, we identify fourteen host-encoded circRNAs that are packaged and focus on the abundant species circESYT2. Expression of HIV-1 Gag alone is sufficient to drive circESYT2 incorporation into virus-like particles, indicating that circRNA incorporation is an intrinsic property of the assembly machinery. Proximity-labeling, mass spectrometry and network analysis of the proteins reveal that circESYT2 is embedded in a filamentous cytoskeleton and interacts predominantly with actin, and functional perturbation in a T-cell line shows that circESYT2 depletion destabilizes actin assembly, reduces actin incorporation into virions and enhances viral egress. Extending this analysis to other retroviruses demonstrates that circESYT2 is more efficiently packaged by murine leukaemia virus than by HIV-1, whereas foamy virus excludes it, indicating virus-specific selectivity in circRNAs capture. These findings suggest circRNAs as bona fide components of retroviral particles and uncover a host RNA–cytoskeleton axis in late replication as an unanticipated layer to host–virus crosstalk.

## Introduction

HIV-1 has a genome of two RNA copies. The virus selectively packages these RNAs using its RNA packaging signal (ψ), ensuring proper viral assembly despite the abundance of host cell RNA (Comas-Garcia et al., n.d.). Computational studies of the 5′ region of HIV-1 RNA, including part of the ψ element, identified secondary structures with loops that facilitate RNA dimerization (Abbink & Berkhout, 2003). Further Nuclear Magnetic Resonance (NMR)-based characterization of a 155-nucleotide region also identified discrete loop structures within ψ that independently direct genomic RNA packaging (Keane et al., 2015). A study demonstrated that the fate of HIV-1 transcripts is decided by a twinned transcriptional start site (TTSS) mechanism. The structural plasticity of HIV-1 RNA in the 5′ leader sequence produces 1G or 3G RNAs with distinct 5′ structures that determine their fate as genomic RNA or mRNA. The 5′ capped RNAs, with 1G, are preferentially packaged in virions as genomic RNA, whereas RNAs with 3G are retained in the cytoplasm as mRNAs (Yasin et al., 2024).

Although the ψ element is critical for selective viral RNA encapsidation, the Gag polyprotein alone exhibits the capacity to package diverse RNAs. Comparative analyses of RNA packaging with or without packageable viral RNA (Ψ⁺/Ψ⁻) have shown that Gag efficiently packages RNA under both conditions. Notably, in the absence of packageable viral RNA, Ψ⁻ particles primarily encapsidate host mRNAs (Rulli et al., 2007). Furthermore, Gag alone is sufficient to assemble virus-like particles (VLPs) in mammalian cells (Jouvenet et al., 2009), with its nucleocapsid domain being essential for multimerization and genome packaging (Berkowitz et al., 1995). Diving deeper into the gag and viral RNA binding using CLIP-seq reveals that in cells, not only nucleocapsid but full-length gag, and subdomains of capsid and nucleocapsid bind to Ψ of the viral RNA for packaging (Lei et al., 2023). Mutational analysis of the HIV-1 Gag protein revealed that impairing its multimerization or membrane association functions disrupts viral RNA packaging, even though particle assembly remains intact (Duchon et al., 2021).

Although extensive studies have elucidated aspects of retroviral RNA packaging, the process remains incompletely understood. RNA packageome analyses have revealed that other retroviruses, such as MLV (murine leukemia virus), encapsidate noncoding RNAs and endogenous retrovirus-derived transcripts (Eckwahl et al., 2015; Garcia et al., 2009; Onafuwa-Nuga et al., 2005). Similarly, HIV-1 packages diverse host RNAs, including tRNAs (Pavon-Eternod et al., 2010), 5S rRNA, U6 snRNA, and Y RNAs (Houzet et al., 2007).

These findings established that retroviruses incorporate a variety of host cellular RNAs in addition to their genomic RNA. Building upon these observations, we sought to investigate whether circular RNAs (circRNAs) are also incorporated into the HIV-1. CircRNAs, formed through backsplicing, lack free 5′ and 3′ ends, rendering them resistant to exonuclease-mediated degradation (Jeck et al., 2013). Their stability and unique structures raise an intriguing possibility that viruses may package circRNAs to enhance viral pathogenesis, or conversely, that host cells may employ them as antiviral factors.

To explore this hypothesis, we performed long-read nanopore cDNA sequencing of HIV-1 virion-associated RNA to characterize the circRNAs in the packageome. Long-read nanopore sequencing enables the capture of full-length circRNAs, allowing reliable detection of back-splice junction (BSJ)–containing reads, which serve as definitive evidence of circRNA molecules (Bhardwaj et al., 2023; Hou et al., 2023; Papatsirou et al., 2024; Rahimi et al., 2021). Furthermore, converting the viral RNA to cDNA not only stabilizes the sample, given the limited abundance and susceptibility of viral RNA to degradation, but also facilitates the generation of a durable and comprehensive viral cDNA library for subsequent analyses. This approach revealed fourteen host-encoded circRNAs packaged into HIV-1 virions, establishing them as a selective cargo class; the abundant circESYT2 emerged as a prime candidate whose functional characterization uncovered an unanticipated circRNA–actin axis constraining viral egress.

## Methods

### Cell culture

Jurkat TAg (JTAg) (Rosa et al., 2015) cells were cultured in Advanced RPMI 1640 (Gibco, catalogue no. 12633–012), and HEK293T [European Collection of Authenticated Cell Cultures (ECACC)] cells were cultured in Gibco advanced Dulbecco’s modified Eagle’s medium (DMEM) supplemented with 5% certified FBS and 2mM L-Glutamine. All cells were maintained in a humidified incubator with 5% CO_2_ at 37^0^C.

### Virus production and infection

For the production of viruses from HEK293T cells, 8 μg of HIV-1 NL4-3 E- (envelope defective)(Pizzato et al., 2007) along with 2 μg of envelope plasmids encoding either Vesicular stomatitis virus G (VSV G) protein for CD4-independent infectivity assays or JR-FL (Rosa et al., 2015) for encoding native envelope of HIV 1 were transfected with the calcium phosphate co-precipitation method. After 48 hours, the virus-containing supernatant was collected and clarified by spinning at 300g for 5 minutes, followed by filtration through a 0.22 μm syringe filter. These viruses were later quantified with the SGPERT assay (Pizzato et al., 2009).

Transfection was done in JTAg cells by electroporation. 8 μg of NL4-3 E- or 8 μg of NLBN Zsgreen (Envelope defective, Nef defective), along with 5 μg of Nef and 2 μg of JR-FL envelope plasmids, were suspended in 200 μL Opti-MEM. Ten million JTAg cells per condition were harvested in the exponential phase by spinning them at 300g for five minutes. Cells were washed with opti-MEM to remove residual serum and debris, then resuspended in 200 μL opti-MEM. Later, the sample (plasmids and cells) was mixed and placed in a 4 mm gap electroporation cuvette, and pulsed in a square wave pattern at 385 V for 6 milliseconds using an electroporator (BTX, Gemini, Harvard Apparatus). Transfected cells were recovered with warm RPMI containing 20% FBS and resuspended in a 6-well plate with 5 ml RPMI containing 5% FBS. Viruses were collected 48 hrs later, filtered, quantified, and used for target cell infection.

### PBMC culture and HIV-1 particle production

Human Peripheral Blood Mononuclear Cells (PBMCs, Product code CL010) were purchased from HiMedia. PBMCs were maintained in Advanced RPMI 1640 supplemented with 5% certified FBS and 2mM L-Glutamine. After thawing, cells were stimulated with 5 μg/mL phytohemagglutinin (PHA) and 50 IU/mL recombinant human IL-2 for 6 hours. The cells were then washed and maintained in IL-2-supplemented media.

For virus production from PBMCs by infection, transiently VSVG pseudotyped NL4-3 was produced by transfecting HEK293T cells. After determining the virus titre with SGPERT assay, PBMC’s were infected at 5 MOI (multiplicity of infection). The medium was changed after 6 hours, and the virus collection from the infected cells was done after 48 hours. The collected viruses were filtered, concentrated and lysed in Trizol for RNA isolation. Similarly, the cells were also lysed in Trizol for RNA isolation.

### Virus production for checking circESYT2 Packaging in other retroviruses

For virus production, HEK293T cells were transiently transfected using the calcium phosphate method. Eight micrograms of NL4-3 E- and 2 μg of JR-FL envelope plasmids for HIV1 particle production, 8 μg of NCA ZSgreen (MLV reporter plasmid) and 2 μg of Ecotropic MLV envelope plasmids for MLV, and pCIES (0.736 μg), pCIPS (1.5 μg), pCIGS (11.84 μg), pΔΦ (11.84 μg) for FV production (Ramdas et al., 2021; Trobridge et al., 2002) were used. Forty-eight hours after transfection, the cell supernatants were collected, filtered, and concentrated. Viral pellets were lysed in TRIzol, and RNA was isolated as described below.

### RNA isolation, cDNA synthesis, and qPCR

Cells were lysed in TRIzol, and after the chloroform phase separation, the upper aqueous layer was processed according to the RNA Clean and Concentrator Kit (Zymo Research). First-strand cDNA synthesis was performed using Superscript III (Invitrogen) reverse transcriptase and random hexamer or oligo(dT) (Thermo Fisher Scientific) as primers. TaqMan probe-based quantitative PCR was performed for checking circESYT2, actin, and HIV-1 gag levels, and for other candidates, SYBR Green I-based RT-qPCR was performed using Dream Taq hot-start DNA polymerase in a Bio-Rad CFX96 real-time PCR. Primer and probe sequences are listed in Supplementary Table 1.

### Western blotting

Cell pellets were lysed with RIPA buffer containing a 2x protease inhibitor cocktail and 5 mM TCEP (a reducing agent) and incubated on ice for 10 minutes. The lysate was centrifuged at 14,000 g for 15 minutes at 4°C, after which the clear supernatant was combined with an equal volume of mPAGE® 4X LDS sample buffer. Proteins were resolved on 8% or 12% SDS-PAGE tricine gels and transferred onto PVDF membranes by electroblotting.

Membranes were blocked with a commercial blocking buffer (Bio-Rad) for 15 minutes, followed by a 1-hour incubation at room temperature with the primary antibody. After washing three times with Tris-buffered saline containing 0.1% Tween 20, the membranes were incubated with secondary antibody for 45 minutes. Blots were washed three more times, and signal acquisition was performed on an Odyssey imager system (LI-COR Biosciences).

### Knockdown of CircESYT2 in JTAg by shRNA

Short hairpin RNA (shRNA) sequences targeting the back-splice junction of circESYT2 were designed in-house based on published literature (Moore et al., 2010) and synthesized by Sigma-Aldrich. These oligonucleotides were amplified under the U6 promoter and cloned into the pTZ57R vector. For transient knockdown experiments, the shRNA plasmid was electroporated into JTAg cells for knockdown, and a shGFP plasmid was used as a control. Knockdown efficiency was evaluated 48 hours post-transfection using RT-qPCR.

### Overexpression of circESYT2 and circESYT2 broccoli

To overexpress circESYT2, the circESYT2 sequence was amplified by PCR and cloned into the Addgene pAV-U6+27-Tornado-Broccoli vector (Litke & Jaffrey, 2019) with the Not I and Sac II sites for infectivity experiments. For imaging studies, the pAV-U6+27-Tornado-Broccoli vector was linearized with Not I and blunt-ended using an End Repair kit, after which the full-length, blunt-ended circESYT2 PCR product was cloned directly upstream of the Broccoli aptamer sequence.

### Opti-prep density gradient centrifugation

Following collection, the viral supernatants were clarified by filtration through a 0.22 μm syringe filter. The filtered supernatants were subsequently layered onto a 20% sucrose cushion prepared in 1× PBS and subjected to ultracentrifugation at 25,000 × *g* for 2 hours at 4°C. After centrifugation, the viral pellets were carefully resuspended in 250 μL of 1× PBS. Pelleted viruses were further purified by Opti-Prep density gradient centrifugation to eliminate contaminating vesicles. A continuous iodixanol gradient ranging from 6% to 18% was prepared in 1× PBS, comprising 11 steps with 1.2% increments. A volume of 250 μL of the concentrated virus suspension was carefully layered onto the gradient and centrifuged for 1.5 hours at 250,000 × g at 4°C using an SW41 Ti swinging-bucket rotor. Eleven gradient fractions were collected sequentially from the top and analysed for reverse transcriptase (RT) activity using an SGPERT assay (Dettenhofer & Yu, 1999; Vaillancourt et al., 2021)

### Localization of Broccoli aptamer-tagged CircESYT2

For performing confocal microscopy, HEK293T cells were seeded onto poly-L-lysine-coated coverslips in 24-well plates and transfected via the calcium phosphate method with relevant plasmids: pcDNA3.1(-) and Tornado Broccoli-CircESYT2. Transfected cells were incubated for 48 hours at 37°C before fixation with 2% paraformaldehyde and subsequent washes with 1× PBS. After subsequent triple washes, nuclear staining was performed with Hoechst (1:10,000) and 20 μM DFHBI-1T for Broccoli aptamer visualization (Filonov et al., 2014), followed by incubation at room temperature. Imaging was conducted with a FV3000 confocal microscope using a 100× objective.

### CircESYT2 detection by FISH (Fluorescence in situ hybridization)

To perform circESYT2 hybridization, biotin-labelled DNA probes complementary to the BSJ of circESYT2 were procured from Sigma. JTAg T cells were then seeded onto poly-L-lysine coated coverslips. The cells were fixed with 2% paraformaldehyde and subsequently permeabilized using 0.1% Triton X-100, followed by blocking with BSA. Next, the permeabilized cells were incubated overnight at 37°C in a dark, humidified chamber with a hybridization buffer containing the biotin probe. The following day, coverslips were washed with 1X PBS and 4X SSC. After washing, the hybridized coverslips were incubated with a 1:500 dilution of streptavidin 488 for two hours, after which they were washed again with 1X PBS containing 0.1% Tween 20. Finally, nuclear staining was conducted using a 1:10,000 dilution of Hoechst. The coverslips were then mounted onto slides, and visualization was performed with an FV3000 confocal microscope using a 100× objective (Ramdas & Chande, 2023)

### Photoactivatable Ribonucleoside-Enhanced Crosslinking and Immunoprecipitation (PAR-CLIP)

PAR-CLIP was performed as previously described (Bhardwaj et al., 2023; Spitzer et al., 2014). Briefly, HEK293T cells were seeded in three 10 cm dishes per condition one day prior to labeling. Twelve hours later, cells were supplemented with 100 μM 4-thiouridine (4SU) and incubated for an additional 16 hours at 37°C. Following incubation, the culture supernatant was removed, and cells were rinsed with ice-cold 1× PBS. Subsequently, cells were subjected to ultraviolet crosslinking with 365 nm light at an energy dose of 0.15 J/cm². Cells were collected by gentle scraping and pelleted by centrifugation at 4°C. The resulting pellet was snap-frozen in liquid nitrogen and stored at −80°C until further use.

For immunoprecipitation, Actin mouse monoclonal antibody (1 μg) or mouse IgG (1 μg; negative control) was incubated with protein G beads for 1 hour to allow coating. Beads were washed three times with citrate phosphate buffer to remove unbound antibodies, then blocked with bovine serum albumin (BSA) and yeast tRNA for 1 hour. Cell pellets were lysed in NP40 lysis buffer, and lysates were clarified by centrifugation at 4°C. The clarified lysate was added to the antibody-coated beads and incubated on a rotator mixer for 5–6 hours at 4°C. Following binding, beads were washed three times each with IP wash buffer and high salt buffer. Beads were then resuspended in 100 μL citrate phosphate buffer.

For downstream analysis, 30% of the bead suspension was subjected to western blot analysis by boiling in an equal volume of 4× Laemmli buffer at 95°C for 5 minutes. The remaining 70% of the sample was processed for RNA isolation using TRIzol after proteinase K digestion.

### CARPID (CRISPR-assisted RNA-protein interaction detection) and Mass spectrometry

Guide RNAs targeting the circESYT2 backsplice junction (BSJ) were designed as described in Bhardwaj et al. and cloned into the guide RNA expressing vector (Addgene #109054). The CARPID BASU-dCasRx plasmid (Addgene #153209) and the guide RNA expressing plasmid were co-transfected into HEK293T cells at a 1:2 ratio. After 48 hours, the culture medium was replaced with medium supplemented with 200 μM biotin and incubated for an additional 15 minutes (Yi et al., 2020). Following incubation, cells were washed three times with ice-cold 1X PBS and harvested.

Cell lysates were prepared using lysis buffer and centrifuged at 14,000 g for 15 minutes at 4°C to remove debris. Protein concentrations in the supernatants were determined using the Bradford assay. Biotinylated proteins were captured by incubating the lysates with MyOne T1 streptavidin beads (Thermo Fisher Scientific). Bound proteins were eluted by boiling the beads in 4X mPAGE® LDS Sample Buffer at 95°C for 5 minutes. The eluted samples were then resolved on an 8% SDS-PAGE gel.

After electrophoresis, the gel was rinsed with MS-grade water and silver stained to visualize protein bands. Differentially expressed protein bands were excised from the gel and subjected to in-gel digestion prior to LC-MS analysis using a TripleTOF system. The acquired spectra were analyzed with Mascot software for protein identification. Further, to investigate the potential molecular interactions associated with circESYT2, we constructed a high-confidence protein–protein interaction (PPI) network using STRING v12.0. We filtered interactions using a stringent combined-score cutoff (>700), representing high-confidence associations. The list of interacting protein pairs obtained from STRING, along with their respective combined scores, was used to generate the final PPI network. In addition to it, to understand the biological pathways significantly associated with the identified proteins (UniProt accessions) from mass spectrometry were mapped to gene IDs, followed by Gene Ontology and pathway enrichment analyses using clusterProfiler and ReactomePA, with significant terms identified by adjusted p-values (FDR < 0.05) and visualized as bar plots.

### Actin staining with Phalloidin 568

JTAg cells were subjected to a shRNA knockdown using the pscalps zsgreen plasmid, which was cloned with a circESYT2 shRNA cassette, along with a shGFP control. Forty-eight hours post-transfection, the cells were seeded onto poly-L-lysine-coated coverslips and fixed with 2% paraformaldehyde. After washing the coverslips with 1X PBS, the cells were permeabilized using 0.1% Triton X-100 and blocked with blocking buffer (1X PBS, 1% BSA, and azide) for one hour. Following the blocking step, cells were stained with Alexa Fluor 568 Phalloidin (Thermo Fisher) as per the manufacturer’s protocol for one hour. After the incubation, the cells were washed three times with blocking buffer, stained with Hoechst (1:10,000), and then mounted on glass slides for confocal imaging.

### Nanopore cDNA library preparation

#### Viral RNA isolation

For isolating Viral RNA from two biological replicates, concentrated viral pellets were lysed in TRIzol. Further, to get sufficient RNA recovery, after the chloroform phase separation, the upper aqueous layer was carefully separated and precipitated using absolute ethanol and glycogen. Further processing was performed using the RNA Clean and Concentrator Kit. To avoid any possible plasmid DNA contamination, on-column DNase I treatment was done. Viral RNA was quantified using a Qubit 4.0 fluorometer with a Qubit HS RNA kit. Further, for nanopore direct cDNA library preparation and sequencing, the SQK-DCS109 kit protocol was followed.

#### Nanopore data acquisition and circRNA analysis

Libraries were prepared with SQK-DCS109 and sequenced on a MinION using a FLO-MIN106 flow cell (ID FAW78155). The run was performed in MinKNOW 3.6.5 (Bream 4.3.16) with on-device basecalling using Guppy 3.2.10, high-accuracy model. Reads with mean Q ≥ 7 (basecaller *min_qscore*) were marked pass and used for downstream analyses.

Raw ONT cDNA FASTQ files were processed with the circNick-LRS workflow (version [2.1]) (Rahimi et al., 2021), isoCIRC (version [1.0.7]) (Xin et al., 2021) and CIRI-long (version [2.0]) (Hou et al., 2023) for detecting circRNAs.

### Assignment of human, viral reads, and RNA Biotype classification

Our pipeline incorporated a multi-step alignment strategy to ensure accurate assignment of sequencing reads. We constructed a combined reference genome comprising the human GRCh38 assembly and the NLBN zsGreen viral sequence, ensuring comprehensive coverage of both host and viral genetic elements. Reads were aligned using minimap2 (v2.24) with splice-aware parameters. The splice-aware configuration ensured accurate mapping of both host mRNA splicing variants and viral transcripts.

In an attempt to quantify the abundance of host linear RNA species in the viral packageome, Nanopore basecalled reads were aligned to the Homo sapiens reference genome (GRCh38, Gencode v44 annotation) using minimap2 with parameters optimized for spliced alignment. Gene-level read counts were then derived using featureCounts (Liao et al., 2014) in long-read mode.

To facilitate further biotype-level analysis, the generated raw counts were combined with gene annotation metadata acquired from Ensembl BioMart (Smedley et al., 2015). For normalization, only genes with detectable read counts across replicates and valid Ensembl IDs were kept.

Transcript abundance was expressed as Transcripts Per Million (TPM), calculated independently for each replicate as described by Wagner et al.(Wagner et al., 2012).

The TPM values from both replicates were averaged to obtain mean TPM per gene, representing the relative expression level of each transcript in the viral packageome. Genes were ranked according to mean TPM to identify the most abundantly packaged host RNAs. The top 1000 expressed genes were used for comparative and visualization analyses in the correlation plot and biotype composition summaries.

The primer, probes, shRNA, and other oligos used are listed in Supplementary Table S1. All the Plasmids, antibodies, and reagents used in this study are listed in the supplementary table S2.

### Statistical analysis

All the statistical analyses were performed in GraphPad Prism 9.1.0. Experiments were performed in three biological replicates, and the data were represented as the means and ± SD of technical replicates unless otherwise stated in the methods. Data were checked for Gaussian (normal) distribution using the Shapiro-Wilk test before performing the statistical analysis. A two-tailed unpaired Student’s t-test was performed to compare two groups. For multiple comparisons, one-way ANOVA (analysis of variance) followed by Dunnett’s or two-way ANOVA followed by Sidak’s post hoc test was performed. All reported differences were ns=non-significant, \**P* < 0.05, \*\**P* < 0.01, \*\*\**P* < 0.001 and \*\*\*\**P* < 0.0001

## Results

### Nanopore sequencing of HIV-1 virion packaged RNA reveals distinct RNA biotypes

To investigate the HIV-1 viral RNA packageome, direct cDNA nanopore sequencing was performed using the Oxford Nanopore platform. Viral stocks produced by transfecting Jurkat TAg T cells were used as the source for viral RNA extraction and subsequent cDNA synthesis (Fig. 1A). Because of the typically low concentration and instability of viral RNA, which increases the risk of RNA degradation, direct cDNA sequencing was selected as the preferred method. Informed by prior findings that randomly primed double-stranded cDNA libraries offer superior sequencing coverage compared to poly(A)-selected libraries (Liefting et al., 2021), random hexamer priming was employed for double-stranded cDNA synthesis. The nanopore cDNA library was then prepared utilizing the SQK-DCS109 kit, incorporating slight protocol modifications to specifically enrich for circRNAs during reverse transcription (Fig. 1B). This approach enabled comprehensive profiling of the HIV-1 RNA packageome.

**Fig 1.**
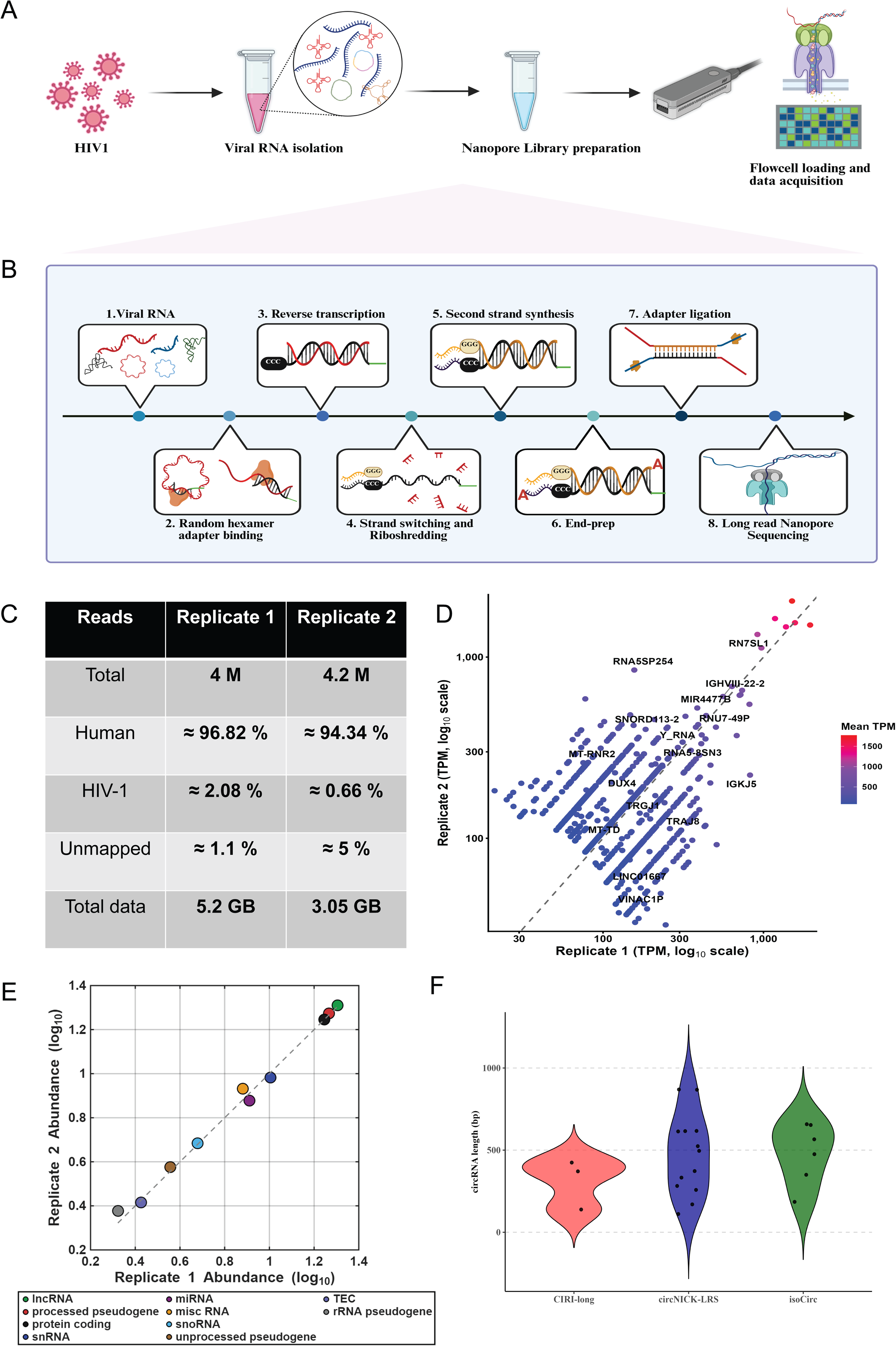
An RNA packageome of HIV-1. Schematics representing **(A)** the flow of viral cDNA sequencing, and **(B)** the Nanopore Direct cDNA sequencing library preparation (SQK-DCS109). **(C)** The mapped reads obtained from two independent biological replicates following high-precision basecalling provide an overview of sequencing consistency and data quality. **(D)** Scatter plot of the top 1000 packaged RNAs in the HIV-1 Packageome. Axes show log-transformed TPM values for replicate 1 and replicate 2. **(E)** Representation of the order of biotype abundance in the Viral Packageome. Each point represents the total abundance (TPM) of a specific RNA biotype detected in HIV-1 viruses across two biological replicates. Axes show log-transformed TPM values for replicate 1 and 2. **(F)** Length distribution of circRNAs detected using CIRI-long, CircNICK-LRS and isoCirc pipelines with default parameters to detect circRNA mature lengths.

To enable comprehensive cDNA synthesis of circRNAs alongside other RNA species, random hexamer adapter primers were used for reverse transcription (/5Phos/ACTTGCCTGTCGCTCTATCTTCNNNNNN-3), which were validated in previous studies (Boldogkői et al., 2018; Della Bartola et al., 2020). Following this step, all subsequent procedures adhered to the standard kit protocol, with further details outlined in the materials and methods section. Viral stock preparation and sequencing experiments were performed in two biological replicates.

Analysis of raw reads from both replicates was conducted using the designated bioinformatics pipeline for evaluation of host and viral transcripts. Among the total reads, 94–96% were aligned to the human transcriptome. This is in congruence with the already done LC/MS analysis of HIV-1 RNA packageome, stating that viral RNA comprises approximately 96% of host RNAs, mainly containing long non-coding RNAs like tRNAs (Šimonová et al., 2019), 0.6–2% aligned to HIV-1 viral sequences, and the remaining 1–5% were unmapped (Fig. 1C). Following normalization by length, the genes with the highest TPM values, like RN7SL1(7SL RNA), Y-RNA and U6 snRNA, were in compliance with the findings of the (Houzet et al., 2007) (Fig. 1D). Further RNA biotype classification following the transcript quantification in the form of TPM revealed that approximately 20–21% of the total TPM corresponded to long non-coding RNAs, 17–18% to protein-coding genes, 18.5 % to the processed pseudogenes with the remainder distributed among small RNAs, intergenic regions, and other biotypes (Fig. 1E). Quality assessment of nanopore sequencing reads was performed using NanoPlot, part of the NanoPack suite, analyzing metrics such as read length, cumulative yield, and average read quality. These results are illustrated in (Fig. S1 A–D) for the first and (Fig. S2 A–D) for the second replicate.

### Structured circRNAs are specifically incorporated into virions

The CircNICK-LRS, isoCirc and CIRI-long pipelines were run on both the biological replicates. Out of the three, circNICK-LRS was the most effective at identifying 14 annotated mature-length circRNAs (Fig. S3A) and 5 unannotated circRNAs. The detected annotated circRNAs were namely CircRBM33, CircNEK1, CircPNN, CircOGA, CircRBM39, CircUBXN4, CircUHRF2, CircGLE1, CircCEP70, CircATAD2B, CircFBXL17, CircERC1, CircMAP7D3 and CircESYT2. Similarly, IsoCirc detected 5 circRNAs namely CircATAD2B, CircSMARCA5, CircFBXL17, CircCCDC88C and CircUBE2Q2. Interestingly, two of the five circRNAs that isoCirc identified were shared by circNICK-LRS. CIRI-long, which performed poorly on our datasets detected only three unannotated circRNAs.(Fig. 1F)

Considering the performance of CircNICK-LRS pipeline, we started with the validation of the 14 circRNA candidates. For the validation of these circRNAs, divergent primers were designed (Supplementary Table 1). Validation was performed by qRT-PCR using these primers to assess circRNA abundance in both virus samples (Fig. 2A) and producer cells (JTAg cells) (Fig. 2B). Viral circRNA quantification was normalized to HIV-1 Gag RNA, while cellular data were normalized to cellular Actin RNA levels. Among the 14 candidates, 8 circRNAs (CircRBM33, CircNEK1, CircPNN, CircOGA, CircRBM39, CircUBXN4, CircUHRF2, and CircESYT2) exhibited greater than 1 log₁₀ higher relative expression, while the remaining six (CircGLE1, CircCEP70, CircATAD2B, CircFBXL17, CircERC1, CircMAP7D3) displayed less than 1 log₁₀ relative expression from viral cDNA. Notably, CircESYT2 expression was particularly robust, showing more than a 10 log₁₀ higher relative expression in viral samples and more than 100 log₁₀ increase in relative expression in T cells. Subsequent validation was extended to circRNAs packaged in viruses produced from JTAg cells, which exhibited more than a 1 log₁₀-fold increase in expression. These were further assessed in wild-type HIV-1 viruses generated from PBMCs (Pooled healthy donors) as well as in producer PBMCs (Fig. 2C and 2D). Remarkably, CircESYT2 demonstrated a more than 30 log₁₀ increase in relative expression in infected PBMCs, correlating with enhanced packaging efficiency in viral particles.

**Fig 2.**
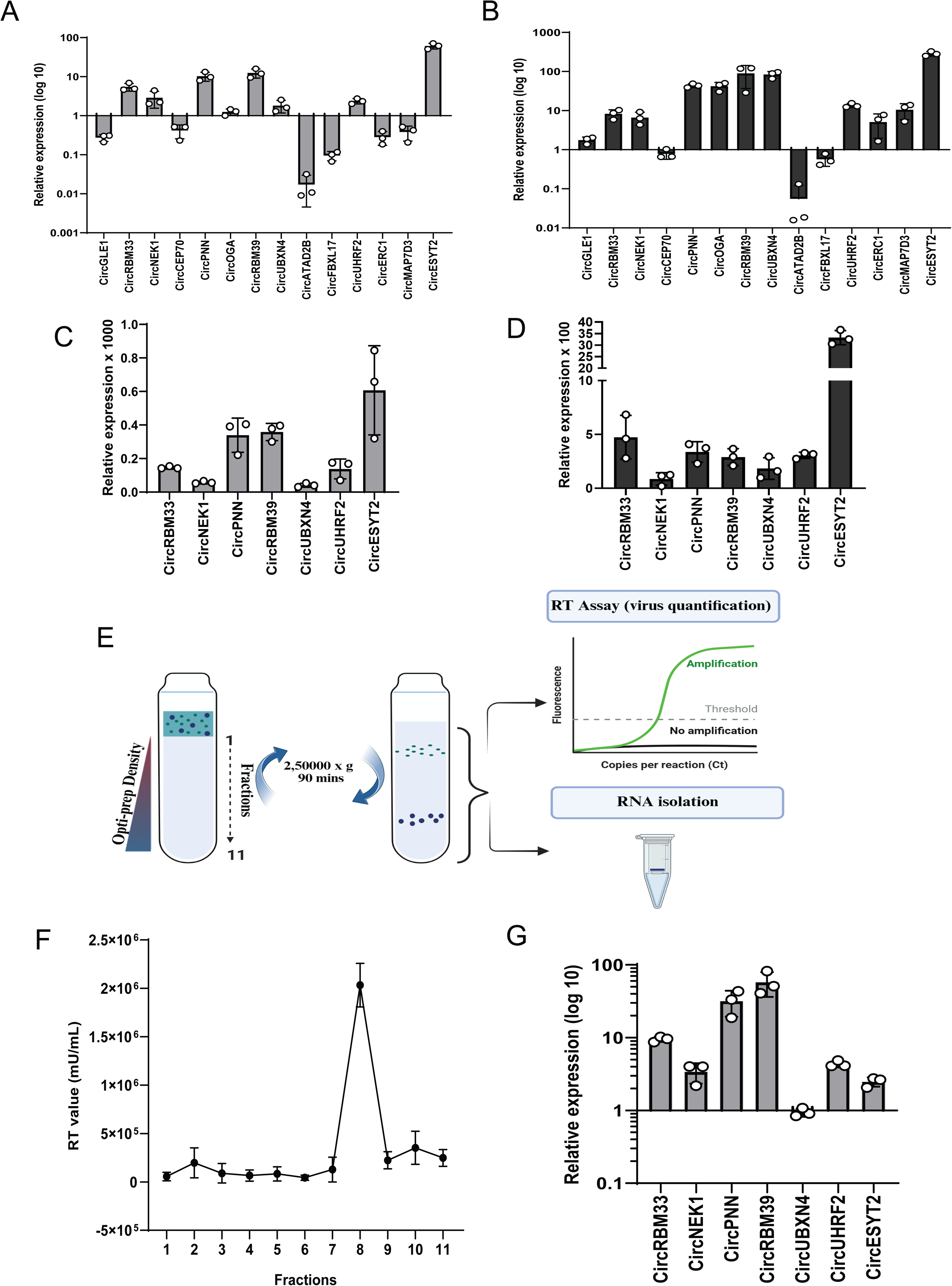
RT-qPCR validation of candidate CircRNAs obtained after nanopore data analysis through the CircNick-LRS pipeline, using cDNA made from RNA of **(A)** HIV-1 viruses produced from Jurkat TAg cells, **(B)** from the producer cells (Jurkat Tag), **(C)** HIV-1 wild-type viruses produced from PBMCs (mixed donors), **(D)** from the producer cells (PBMCs). Data obtained from cells was normalized to Actin, and data obtained from viruses was normalized to HIV-1 Gag. Validation of the circRNAs incorporated in the HIV-1 viruses. **(E)** Opti-prep density gradient centrifugation workflow for separating viruses from vesicles. **(F)** SGPERT assay performed on the gradient fractions to capture virus-containing fractions. **(G)** RT-qPCR validation of the selected circRNA candidates using RNA isolated from the 8^th^ Fraction (from F), showing the highest RT activity. Data was normalized to Gag RNA.

Due to the notably high packaging efficiency of these circRNAs, their specific selection for viral packaging warranted further investigation. To explore this, the free energies of these circRNA candidates were evaluated using the RNAfold webserver (http://rna.tbi.univie.ac.at/cgi-bin/RNAWebSuite/RNAfold.cgi), which revealed that these molecules exhibit particularly low predicted free energies Fig. S3B. This implies greater structural stability, since for any given RNA length, lower free energy corresponds to a more stable RNA fold (Workman & Krogh, 1999). Enhanced stability may thus underlie their specific inclusion in viral particles. Furthermore, we also validated the BSJs of these 7 circRNAs, amplified from viruses using Sanger sequencing, and confirmed the sequences Fig. S3C-H, Fig. 3A.

**Fig 3.**
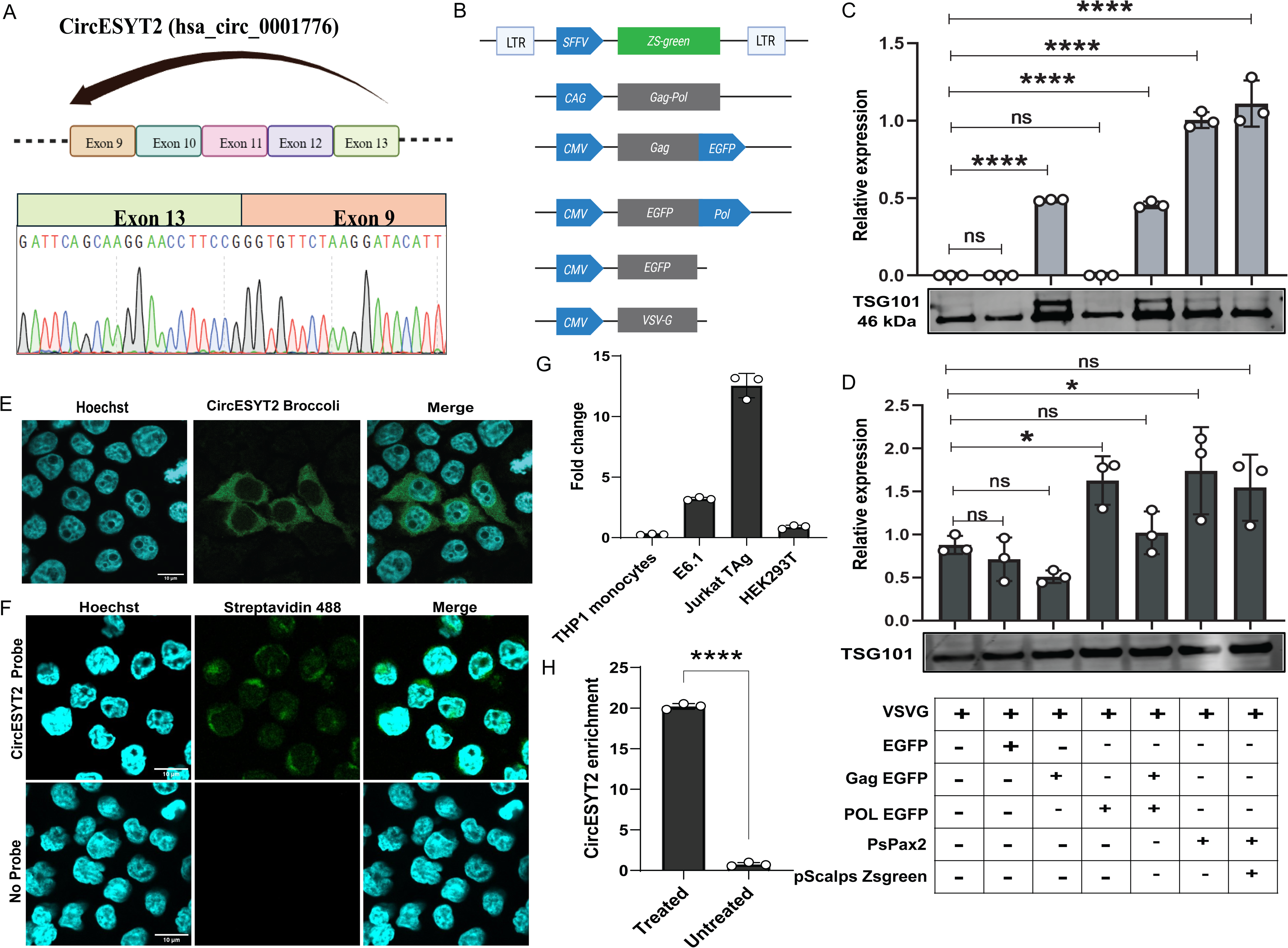
HIV-1 Gag selectively incorporates circESYT2 in the vesicles. **(A)** Sanger sequencing confirmation of the backsplice junction of circESYT2 (hsa_circ_0001776). **(B)** Schematics of the plasmids used for dissecting the vesicle packaging specificity of circESYT2. circESYT2 RT-qPCR from RNA isolated from **(C)** Vesicles and **(D)** the Producer cells (HEK293T). TSG101 is taken as a marker for vesicles. Cellular CircESYT2 data were normalized to actin. Confocal imaging of HEK293T cells showing **(E)** CircESYT2 Broccoli (green) localization in HEK293T cells. **(F)** CircESYT2 FISH in JTAg cells and visualization with streptavidin 488. **(G)** CircESYT2 expression checking in the cell lines. **(H)** RNAseR resistance checking for circESYT2 using TaqMan RT-qPCR. One-way ANOVA with Dunnett’s multiple comparisons test to assess the significance between the multiple groups (C and D). The unpaired two-tailed Student’s t-test was used to assess the significance between two groups (H). (n = 3; ± SD), ns=not significant.

To further strengthen our findings on the packaging of circRNAs in viruses, we further purified the viral suspension. The viruses were concentrated on a 20% sucrose cushion; however, some microvesicles, similar in size to the viruses, might still contaminate the viral stocks. Therefore, we separated the concentrated viral stocks from vesicles using a 6-18% continuous iodixanol gradient (OptiPrep), as shown in Fig. 2E. The 11 gradients collected were analyzed for viral RT activity using the SGPERT assay. Among the 11 gradient fractions, the highest RT activity was detected from the 8th fraction (Fig. 2F). The RNA isolation from the 8th fraction was performed according to the viral RNA isolation protocol described in the Methods. RTPCR screening of these seven candidates revealed that, except for CircUBXN4, all other candidates exhibited relatively more than 2 log₁₀ higher expression, indicating their presence in the viruses and suggesting virion-specific incorporation (Fig. 2G).

### Gag is required for the incorporation of circESYT2 in the virions

Given its notable abundance in T cells, primary cells, and virions, circESYT2 is well-suited as a candidate for investigating the mechanisms underlying viral circRNA packaging. CircESYT2 (hsa_circ_0001776) is a 495 bp circRNA produced by back splicing of exon 13 to exon 9 of the ESYT2 gene (Extended Synaptotagmin 2), as illustrated in Fig. 3A. HIV gag, a structural protein crucial for viral particle assembly, is also recognized for its role in RNA binding and determining selective encapsidation of genomic and non-coding RNAs in virions.

To assess whether gag alone is sufficient for circESYT2 packaging, a series of plasmid constructs was generated (Fig. 3B), and VSVG vesicles were produced for functional studies. Analysis of vesicular RNA revealed that the gag protein, in isolation, can mediate circESYT2 incorporation into vesicles. However, this packaging was significantly enhanced—approximately doubled—in vesicles containing the gag-pol polyprotein or in authentic virions (Fig. 3C), suggesting virus-specific mechanisms are at play in circESYT2 packaging. TSG101 served as a marker for vesicles during these experiments. Importantly, other vesicle types carrying cargoes—such as VSVG only, EGFP-VSVG, and EGFPPol-VSVG—failed to package circESYT2, even though the circRNA was present in the producer cells (Fig. 3D), supporting the specificity of circESYT2 packaging mediated by gag and viral context.

### CircESYT2 shows uniform cytoplasmic localization

Given the direct packaging of circESYT2 into viruses, we were curious about its subcellular localization. To visualize circESYT2, it was tagged with a Broccoli aptamer and analyzed by confocal microscopy in HEK293T cells. After performing confocal microscopy, circESYT2 showed uniform cytoplasmic distribution (Fig. 3E). As this was the localization of the overexpressed circESYT2, we also wanted to validate the localization of the endogenous circESYT2. To this end, we performed FISH for circESYT2 in JTAg cells. Imaging the cells after probing with streptavidin 488 revealed cytoplasmic localization (Fig. 3F), validating our previous results.

### Specificity of circESYT2 expression and retroviral packaging

We assessed the RNase R resistance of circESYT2 using RNA extracted from JTAg cells and observed an approximately 20-fold enrichment of circESYT2 in RNase R-treated samples compared to untreated controls, with normalization to GAPDH (Fig. 3H). Following confirmation of circESYT2’s circular nature, we quantified its expression levels across several cell types, including monocytes, Jurkat E6.1 T cells, JTAg T cells, and HEK293T cells. JTAg T cells exhibited the highest circESYT2 expression, approximately 10-fold higher than that of other cell lines (Fig. 3G).

To determine whether circESYT2 packaging is virus-specific, we analyzed its incorporation into different retroviruses: HIV-1, MLV, and FV. Notably, MLV packaged circESYT2 at levels approximately 3000-fold higher than HIV-1, whereas FV showed minimal or no packaging (Fig. S4A), indicating preferential incorporation by specific retroviruses.

Subsequently, we modulated circESYT2 levels by knockdown or overexpression to investigate its impact on viral replication kinetics. Two shRNAs targeting circESYT2 were designed and tested in HEK293T cells, with shRNA_1 achieving a knockdown efficiency reducing circESYT2 levels to less than 25% without affecting ESYT2 mRNA expression (Fig. S4B, S4C). Importantly, knockdown in producer cells corresponded with a near-complete loss of circESYT2 packaging into virions (Fig. S4D). Similarly, knockdown in JTAg cells with shRNA_1 resulted in around a 60 % loss of circESYT2 expression in JTAg cells (Fig. S4E), without affecting mRNA and protein expression (Fig. S4F)

For overexpression, circESYT2 was expressed using a ribozyme-based Tornado vector. Overexpression in JTAg T cells resulted in a 2–5-fold increase in circESYT2 levels (Fig. S4G), whereas HEK293T cells exhibited an approximately 800-fold increase compared to empty vector controls (Fig. S4H). This elevated expression translated into a 2000–4000-fold enhancement of circESYT2 packaging in viruses produced from overexpressing cells (Fig. S4I). These data demonstrate that circESYT2 expression and packaging can be robustly modulated in a cell- and virus-specific manner.

### CircESYT2 interacts with cellular actin, facilitating its packaging into virions

Since Gag is not the sole protein involved in circESYT2 packaging, we sought to identify other potential interactors of this circRNA. To do this, we employed CARPID, which uses a catalytically dead CasRx fused to a biotin ligase. We directed the CasRx to circESYT2 using a guide RNA targeting its backsplice junction (BSJ), while a guide RNA against the luciferase gene served as a negative control. Further, the pull-down of biotinylated proteins was performed according to the methods.

Following SDS PAGE run and silver staining of the gel (Fig. 4A), we observed two distinct protein bands uniquely present in the circESYT2 guide RNA sample. Mass spectrometry analysis identified these bands as Actin (ACTA, ACTB) (Fig. 4B) and Tubulin (TUBA1A) (Fig. 4C), based on their protein scores and expected molecular weights. Additionally, according to our PPI network analysis, actin was portrayed as having high confidence as a central regulator of the network (Fig. S5A). The GO term “structural constituent of cytoskeleton” is overrepresented in our GO-MF analysis, and terms like “Phagosome” and “motor proteins” are significantly enriched in the KEGG-based pathway analysis, emphasizing that circESYT2 is engaging with the cytoskeletal proteins (Fig. S5B and S5C).

**Fig 4.**
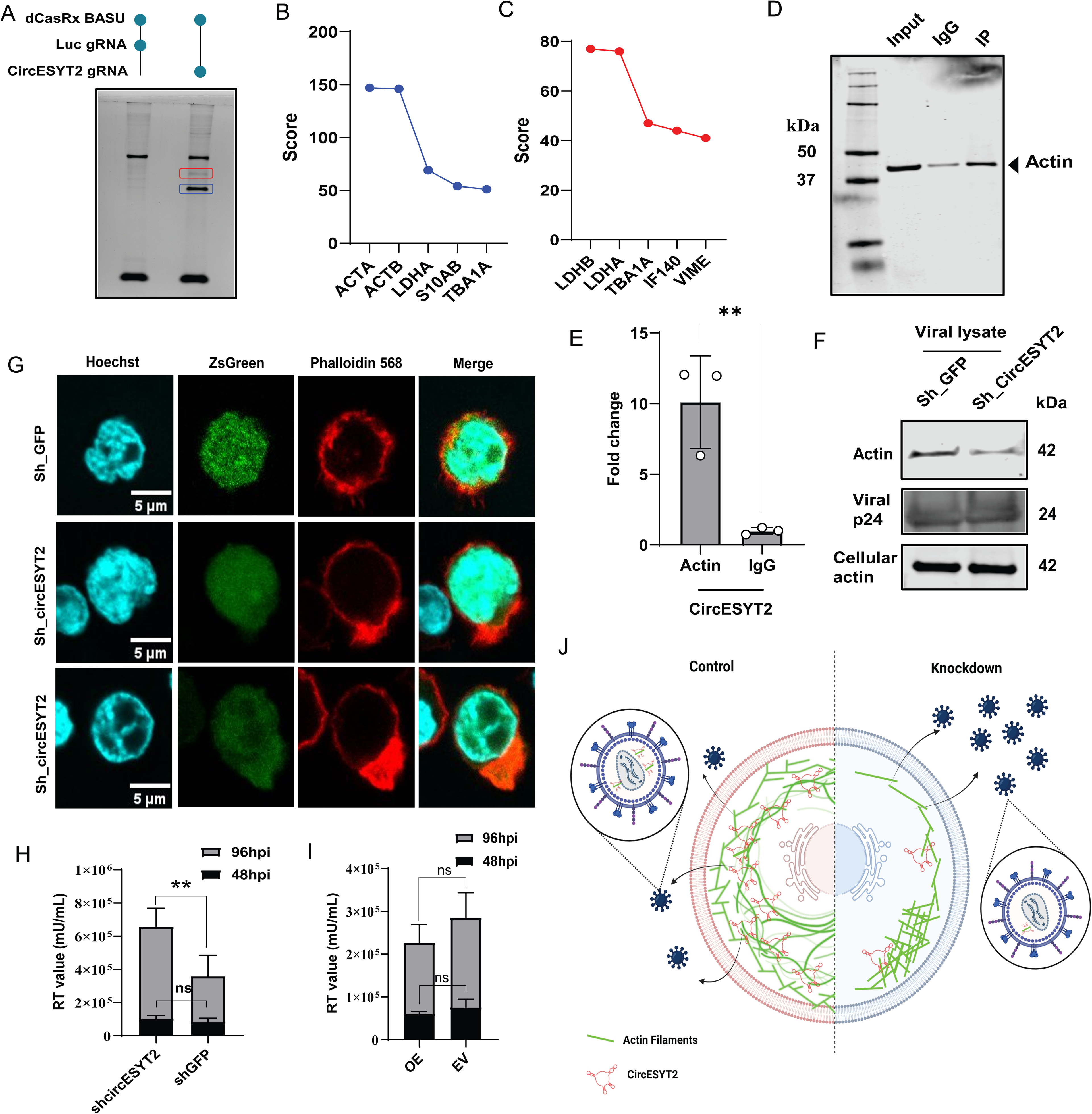
CircESYT2 binds to actin, stabilizing its cytoskeletal assembly **(A)** Silver-stained SDS PAGE showing the differentially enriched biotinylated protein bands. Protein bands analyzed with mass spectrometry are highlighted. The score obtained from **(B)** the blue-highlighted band and **(C)** the red-highlighted band. Actin PARCLIP **(D)** Western blot showing enrichment of actin in pulldown samples as compared to IgG. **(E)** RT-qPCR showing enrichment of circESYT2 in pulldown sample. **(F)** Western blot of viral lysates showing levels of actin protein packaged in virions in circESYT2 knockdown and control conditions. Viral p24 and cellular actin depict equal amounts of virus and cells, respectively. **(G)** Confocal images of phalloidin 568-stained circESYT2 knockdown and control cells. Scale bar represents 5µm. RT assay showing the HIV-1 wild type viral egress from the **(H)** circESYT2 knockdown and control **(I)** circESYT2 overexpression (OE) and Empty vector (EV) JTAg cells after 48- and 96-hours post-infection (hpi). **(J)** Schematics depicting the overall summary of the effects of CircESYT2 altered levels on actin organization and virus egress. For Fig. E, the unpaired two-tailed Student’s t-test was used to assess the significance between the two groups. (n = 3; ± SD), ns= not significant. For Figures H and I, a Two-Way ANOVA with Sidak’s multiple comparisons test was used to analyze the grouped data.

Previous studies have demonstrated that cellular Actin is packaged into HIV-1 virions (Ott et al., 1996), and in our CARPID analysis, Actin was highly enriched in association with circESYT2. To investigate whether circESYT2 directly interacts with Actin for packaging, we performed PAR-CLIP to confirm the interaction. The pull-down was confirmed for Actin protein enrichment (Fig. 4D). RT-qPCR analysis of Actin-crosslinked RNA revealed over a 10-fold enrichment of circESYT2 compared to the IgG control (Fig. 4E). These results confirm that Actin directly binds to circESYT2.

### Actin Protein assembly and packaging in virions is dependent on CircESYT2

If Actin binds to circESYT2 and is packaged into viruses, then knocking down circESYT2 should impair Actin packaging. To test this, viruses were produced under circESYT2 knockdown and control conditions. Virus preparations, normalized by RT activity, were concentrated on a sucrose cushion, and the resulting pellets were lysed in 4X LDS Sample Buffer containing TCEP. Western blot analysis of the viral lysates revealed a reduction in Actin levels in the circESYT2 knockdown viruses compared to controls, while cellular Actin levels remained unchanged (Fig. 4F).

Intrigued by the reduction in viral Actin packaging despite unchanged cellular Actin levels, we investigated whether changes in Actin organization or dynamics could explain this observation. JTAg cells were electroporated with shRNA targeting circESYT2 to achieve knockdown, and both knockdown and control cells were stained with Phalloidin 568 to visualize Actin fibers. Confocal microscopy revealed that control cells exhibited evenly distributed Actin throughout the cytoplasm, whereas circESYT2 knockdown cells showed Actin clustered in distinct regions within the cytoplasm (Fig. 4G).

### Actin remodeling due to the absence of circESYT2 facilitates viral budding

Previous studies have shown that HIV-1 assembly and release sites are characterized by low levels of cortical actin. The virus achieves this reduction in branched actin either by recruiting the host factor Arpin, which inhibits the ARP2/3 complex and promotes actin debranching (Dibsy et al., 2023), or by depolymerizing actin at budding sites through the action of cellular oxidoreductases such as MICAL1 (Serrano et al., 2024).

To investigate the impact of altered circESYT2 levels on viral replication, we infected an equal number of circESYT2 knockdown and overexpressing JTAg T cells with equal reverse transcriptase (RT) units of replication-competent wild-type HIV-1. Viral egress was measured at multiple time points using the SGPERT assay. Within the first 48 hours post-infection, no significant differences in viral egress were observed between knockdown, overexpressing, and control cells. However, at 96 hours, viral egress in circESYT2 knockdown cells was approximately doubled compared to controls (Fig. 4H). circESYT2 overexpression (OE) resulted in a modest reduction of viral egress relative to empty vector (EV) cells (Fig. 4I). These results suggest that actin disassembly due to circESYT2 knockdown improves HIV-1 release from these cells. Based on the results so far, we have summarized our findings in the model represented in (Fig. 4J).

## Discussion

Comprehensive RNA packageome studies across multiple viral systems have significantly advanced our understanding of virion-associated viral and host RNAs. Evidence for host RNA packaging in viruses was first reported in the 1980s, when host tRNAs were identified as major RNA species packaged in vesicular stomatitis virus (VSV) and HIV-1 (Isaac & Keene, 1981; Jiang et al., 1993). DNA viruses such as herpesviruses (Sciortino et al., 2001) and human cytomegalovirus (Terhune et al., 2004) have also been shown to incorporate cellular RNAs into their virions.

Despite extensive studies, one RNA species has remained largely unexplored within viral RNA packageome analyses: circRNAs. In this study, we investigated the incorporation of circRNAs into HIV-1 virions using long-read nanopore cDNA sequencing. Our linear RNA analysis aligned with previous Illumina-based observations, revealing high transcript per million (TPM) counts for 7SL RNA (ONAFUWA-NUGA et al., 2006). Using the circNICK-LRS pipeline, we identified 14 candidate circRNAs, which were subsequently validated by RT-qPCR and confirmed through Sanger sequencing. Minimal free energy (MFE) analysis done using RNAfold webserver (Kazanskii et al., 2024) indicated high structural stability for these circRNAs, consistent with their circular nature.

Interestingly, circRNAs such as CircGLE1, CircMAP7D3, and CircERC1, despite being expressed in T cells, exhibited minimal packaging, suggesting that expression levels alone do not determine their incorporation into viral particles. These findings support the idea of non-random RNA packaging in HIV-1, as also described by Onafuwa et al. for MLV. Further analyses using OptiPrep gradient-purified virions confirmed our hypothesis that circRNA packaging is both selective and regulated. In particular, circESYT2, which showed high expression and significant packaging efficiency, was further characterized. Vesicular studies using VSV-G pseudotyped vesicles demonstrated that the Gag protein is necessary and sufficient for circESYT2 packaging. In contrast to observations in MLV, where Ψ⁺ and Ψ⁻ particles contain comparable RNA levels (Rulli et al., 2007), circESYT2 packaging was almost two times relatively higher in Ψ⁺ particles compared to Ψ⁻ or Gag-only/Gag+Pol particles.

CircESYT2 exhibited elevated expression in T cells compared to other cell types and demonstrated virus-specific packaging preferences, with the highest incorporation observed in MLV virions relative to HIV-1. Proximity ligation assays further revealed circESYT2 as a regulator within the cytoskeletal filament network. Protein-protein interaction analysis via STRING, informed by mass spectrometry data, identified actin as the primary interactor of circESYT2, a finding verified by PAR-CLIP.

Prior studies have established substantial actin incorporation into HIV-1 virions (Rahman et al., 2014). Early research proposed that the HIV-1 Gag nucleocapsid domain binds actin filaments (Liu et al., 1999; Orlova et al., 2014), though subsequent work demonstrated this domain to be dispensable for actin packaging (Stauffer et al., 2014). Our findings extend these observations by implicating circESYT2 in actin regulation during viral assembly. Specifically, circESYT2 knockdown in T cells markedly reduced actin packaging into virions. Confocal imaging further showed disrupted actin organization in knockdown cells, characterized by clustered actin distribution rather than the uniform actin network observed in controls.

Intriguingly, this actin cytoskeletal disruption correlated with enhanced viral particles being detected from supernatants in circESYT2-deficient cells compared to controls, suggesting increased viral egress. Collectively, these results position circESYT2 as a novel mediator of actin dynamics and cytoplasmic organization. Its depletion disassembles actin filaments, thereby alleviating cytoskeletal restraint on virus budding and underscoring the actin network as a barrier to HIV-1 release.

## Acknowledgments

P.M and A.P. received fellowships from UGC and DBT respectively. A.C. is an EMBO Global Investigator at IISERB. This work was supported by the DBT/Wellcome Trust India Alliance Fellowship (IA/I/18/2/504006), and by European Molecular Biology Organization (#5753), Lady Tata Memorial Trust Young Scientist Award, Indian Council of Medical Research (EM/dev//CAR/-2024-01-0113/F2/2024), and Anusandhan National Research Foundation (ANRF) grant # CRG/2023/000720 (to A.C.). The funders had no role in study design, data collection and analysis, decision to publish, or preparation of the manuscript. The authors are thankful to M. Pizzato, the NIH AIDS reagent program, and NIBSC for reagents and cell lines.

## Ethics statement

The Institute Ethics Committee of Indian Institute of Science Education and Research Bhopal approved the study (IISERB/IEC/Certificate/2018-II/04).

## Author contributions

Data curation: P.M., A.P. Software: P.M., A.P., A.C. Formal analysis: P.M., A.P., A.C. Validation: P.M., A.P. Investigation: P.M., A.P., A.C. Methodology: P.M., A.P., A.C. Resources, conceptualization, supervision, and funding acquisition: A.C.

## Supplementary figures

**Fig S1.**
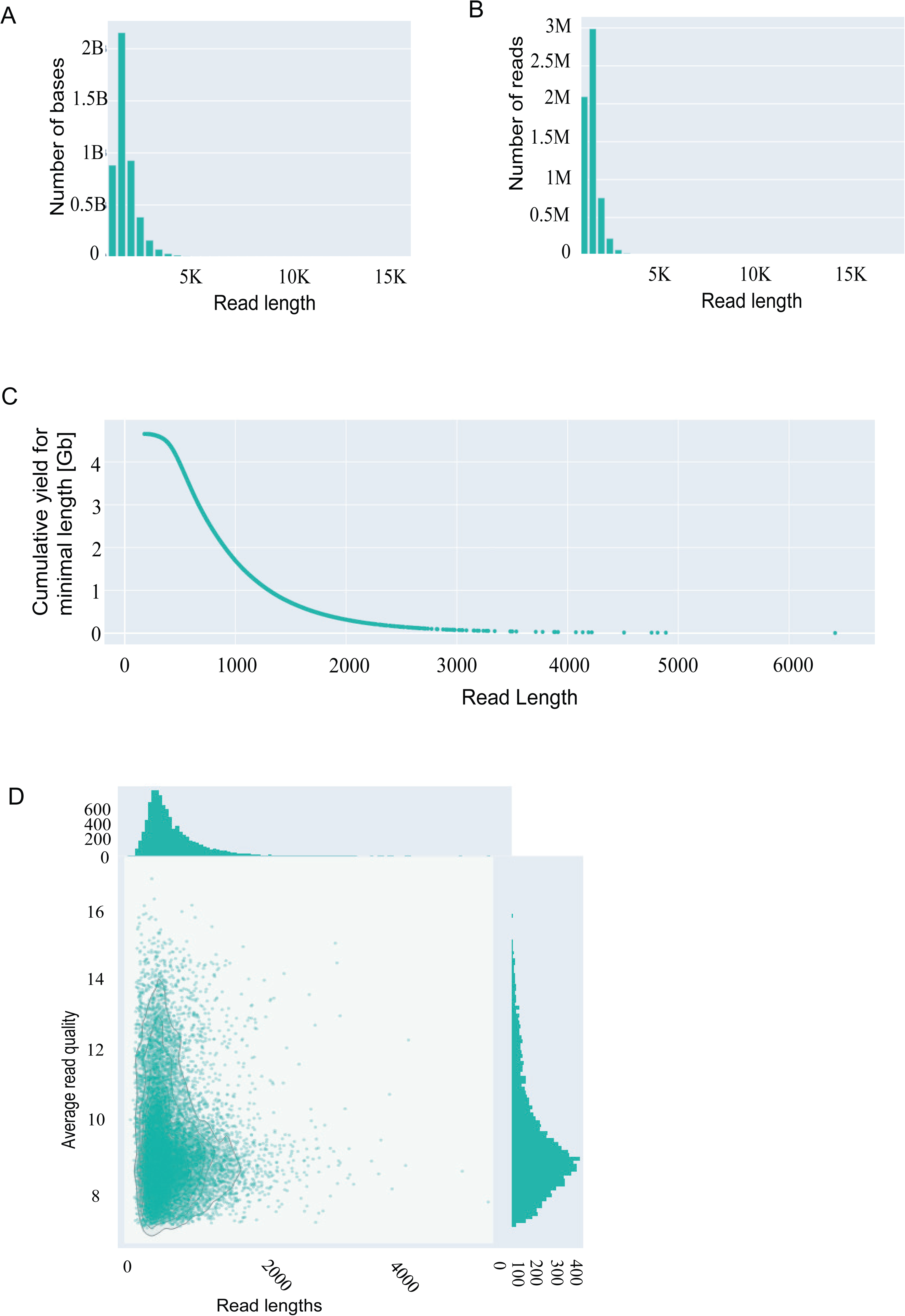
Nanoplot summary metrics of Nanopore HIV-1 cDNA sequencing Replicate 1 **(A)** Number of bases vs. read length, **(B)** number of reads vs. read length, **(C)** cumulative yield for minimal length vs. read length, and **(D)** average read quality vs. read length. These plots summarize read length distribution, sequencing yield, and quality characteristics of the dataset.

**Fig S2.**
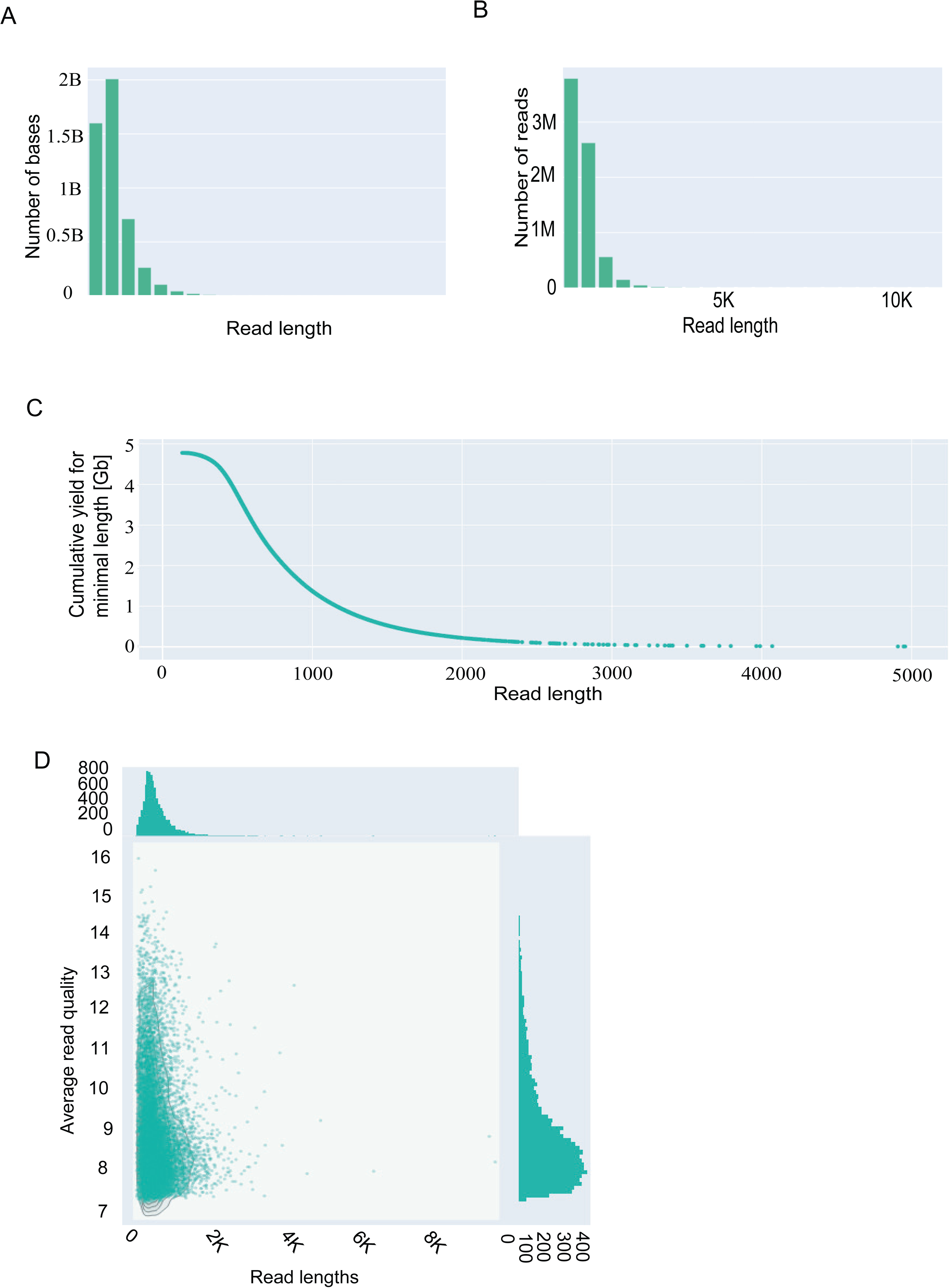
Nanoplot summary metrics of Nanopore HIV-1 cDNA sequencing Replicate 2 **(A)** Number of bases vs. read length, **(B)** number of reads vs. read length, **(C)** cumulative yield for minimal length vs. read length, and **(D)** average read quality vs. read length. These plots summarize read length distribution, sequencing yield, and quality characteristics of the dataset.

**Fig S3.**
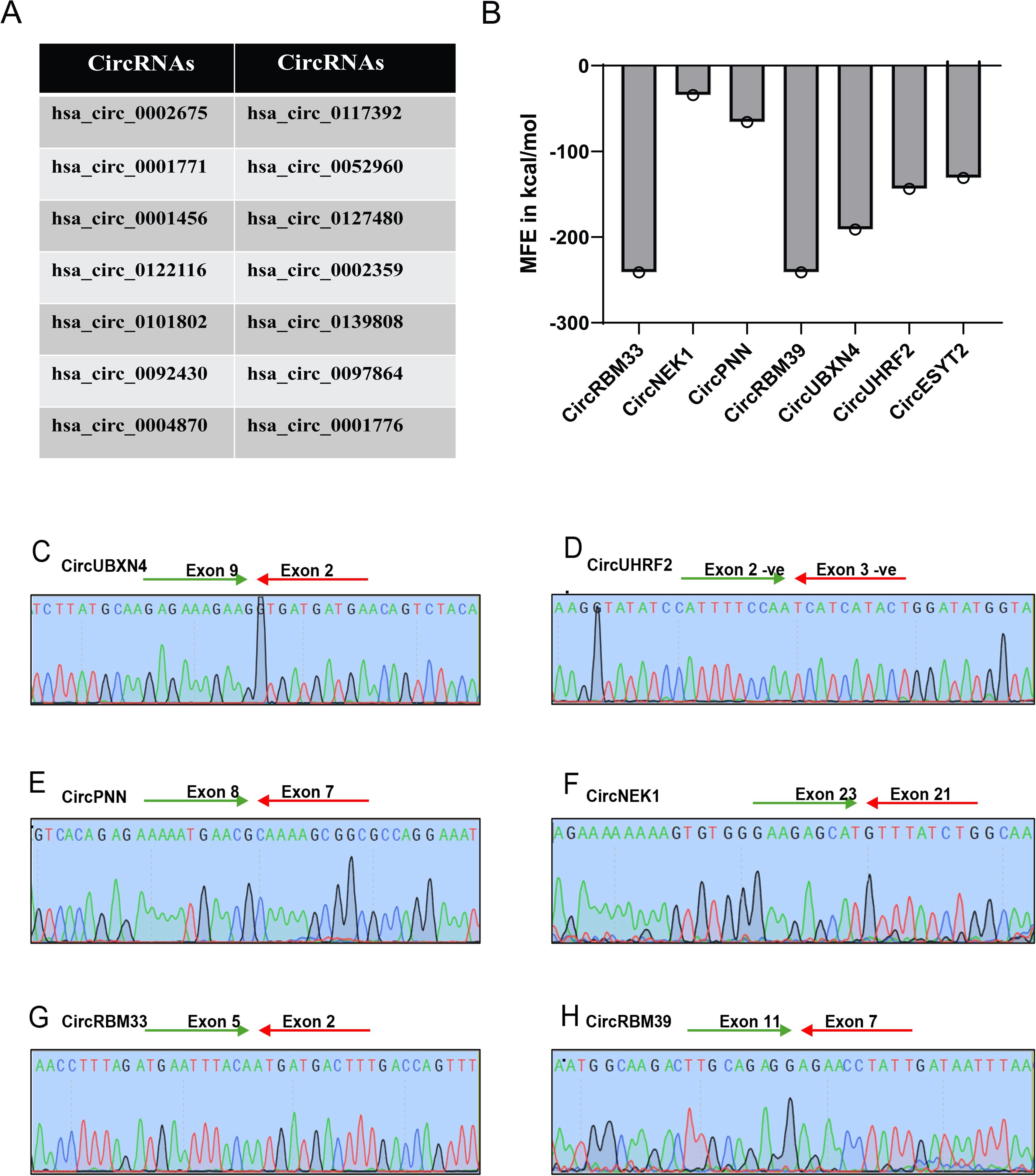
**(A)** 14 Annotated CircRNA candidates identified by circNICK-LRS. **(B)** Minimal free energy (MFE) of validated circRNAs obtained from RNAfold webserver. Sanger sequencing of the back-splice junctions validated from viral cDNA **(C)** CircUBXN4 **(D)** CircUHRF2 (formed from antisense/negative DNA strand) **(E)** CircPNN **(F)** CircNEK1 **(G)** CircRBM33 **(H)** CircRBM39

**Fig S4.**
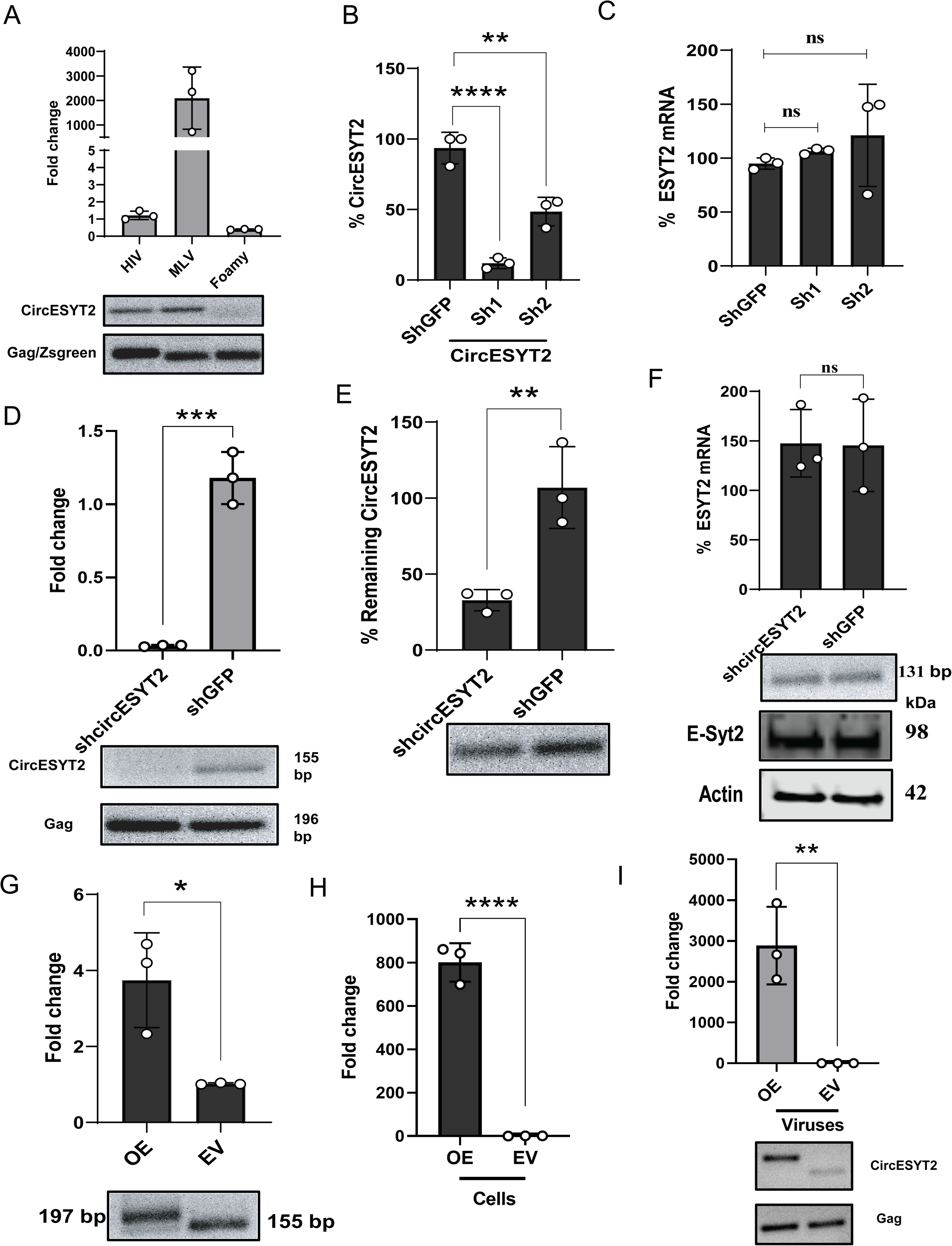
**(A)** RT-qPCR of CircESYT2 packaging checking in other retroviruses, such as MLV and foamy viruses, as compared to HIV-1. Data of HIV-1 and foamy were normalized to their respective Gag RNAs, and MLV was normalized to Zsgreen (NCA Zsgreen). **(B)** Taq-Man RT-qPCR checking of CircESYT2 levels in shRNA knockdown using shcircESYT2_1 (sh1) and shcircESYT2_2 (sh2) using shGFP as control in HEK293T cells. **(C)** RT- qPCR checking of Linear ESYT2 mRNA in circESYT2 knockdown HEK293T cells. **(D)** Taq-Man RT- qPCR checking of circESYT2 in HIV-1 viruses produced from circESYT2 knockdown and control HEK293T cells. The panel below displays the respective gel images for circESYT2 and HIV-1 gag. **(E)** TaqMan RT-qPCR of circESYT2 for checking the shRNA-based knockdown of circESYT2 in JTAg cells, shGFP served as a control. **(F)** RT-qPCR analysis of the linear ESYT2 mRNA levels after the circESYT2 knockdown in JTAg cells. The panel below displays the respective gel image, E-Syt2 Western blot (98 kDa), and actin (42 kDa). **(G)** RT-qPCR verification of circESYT2 overexpression in JTAg cells **(H)** RT-qPCR verification of circESYT2 expression in HEK293T cells overexpressing circESYT2 in comparison to empty vector cells. **(I)** RT-qPCR verification of circESYT2 packaging in HIV-1 viruses produced from HEK293T cells overexpressing circESYT2 in comparison to empty vector cells. The panel below represents the gel images of packaged circESYT2 and HIV-1 gag. The data for B, C, E, F, G, and H were normalized to actin. The data for D and I were normalized to HIV-1 gag. For B and C, the one-way ANOVA with post hoc Dunnett’s multiple comparisons test was performed. For D, E, F, G, H, and I, an Unpaired two-tailed t-test was performed.

**Fig S5.**
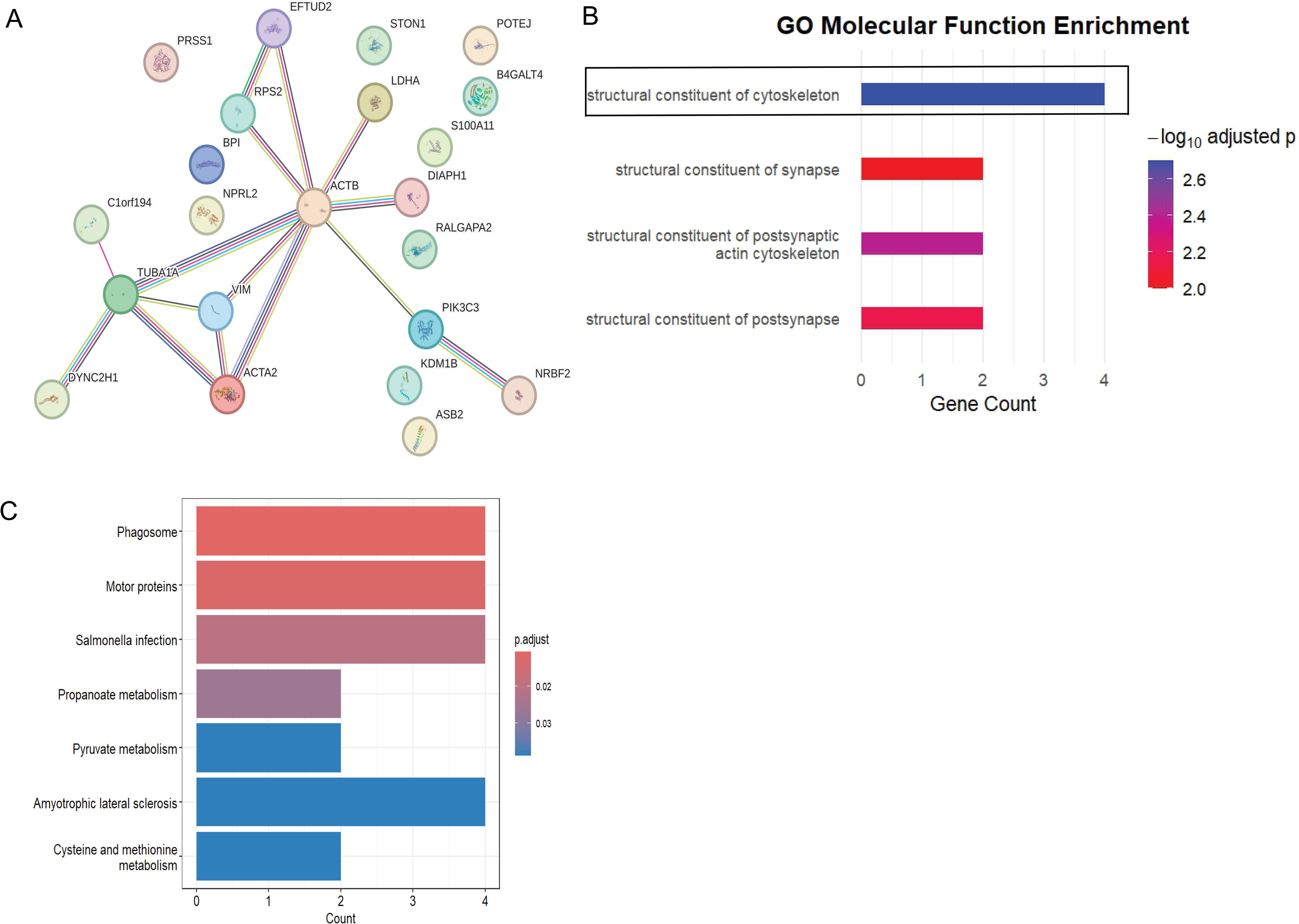
**(A)** High-confidence interactions among the identified proteins were visualized. Nodes represent proteins (UniProt IDs), and edges denote evidence-supported interactions aggregated by the STRING database. **(B)** Gene Ontology (GO) Molecular Function enrichment of circESYT2-associated proteins. Significantly enriched MF terms (FDR< 0.05) identified using clusterProfiler are shown, ranked by gene count. **(C)** KEGG based pathway analysis.

